# PML-driven sumoylation of PML/RARA-bound co-repressors drives immortalization of primary hematopoietic progenitors

**DOI:** 10.64898/2026.01.05.697641

**Authors:** Hsin Chieh Wu, Emmanuel Laplantine, Cécile Esnault, Michiko Niwa-Kawakita, Yi Zhang, Marie-Claude Geoffroy, Hugues de Thé

## Abstract

While acute promyelocytic leukemia (APL) is always driven by fusions involving one of the three retinoic acid receptors, why PML and RARA are the preferred fusion partners has remained largely unsettled. Here we demonstrate that corepressor (NCoR) binding onto the RARA moiety of PML/RARA is required for hematopoietic progenitor immortalization. We then establish that PML-mediated tethering of the UBC9 SUMO conjugating enzyme onto PML/RARA enforces SUMO2 conjugation of multiple RARA partner proteins, notably the NCoR complex, boosting its repressive power. PML mutants that fail to recruit UBC9 yield PML/RARA fusions that neither promote NCoR sumoylation nor transformation. Conversely, direct UBC9/RARA fusion drives both efficient NCoR sumoylation and immortalization. Sumoylation inhibitors re-activate retinoic acid target genes in PML/RARA-, but not RARA-, expressing progenitors and trigger APL differentiation. Thus, fusion of PML to RARA entails an unexpected key gain of function that boosts RARA-mediated transcriptional repression through sumoylation of PML/RARA-bound protein, explaining the recurrent implication of PML and RARA in APL pathogenesis.

## Introduction

Acute promyelocytic leukemia (APL) is driven by fusion proteins always involving one of the three retinoic acid receptors (Geoffroy et al., 2021; Zhang and Qiu, 2025), pointing to the critical role of deregulated retinoic acid (RA) signaling in initiation of the disease. In most patients, these fusions involve RARA and PML (de The et al., 1990). While the pathways involved in retinoic acid (RA) and arsenic trioxide (ATO) triggered PML/RARA degradation and APL cure have been largely deciphered (de The et al., 2017; Dos Santos et al., 2013; Rerolle et al., 2024), why RARA and PML are the preferred fusion partners driving APL initiation remains overall poorly understood. RARA binds corepressors more avidly than other RARs and repression by retinoic acid receptors play an important role in development (Weston et al., 2003). PML/RARA disrupts both PML nuclear body formation and nuclear receptor-mediated transcriptional control (de The et al., 2017). Dominant-negative RARA mutants exert transforming activities in a variety of biological systems, including some breast cancers (Saitou et al., 1995; Tan et al., 2015; Tsai et al., 1992). PML/RARA is an even more potent repressor than RARA and poorly activates most RA-sensitive genes or reporters upon RA binding (de Thé et al., 1991). Yet, the molecular bases for PML/RARA super-repressive activity remain imperfectly understood. PML-facilitated PML/RARA homodimers formation was proposed to enhance HDAC and corepressor binding (Lin and Evans, 2000), while tethering of PML-bound repressors such as Daxx (Zhu et al., 2005) may also contribute to PML/RARA-driven repression. Yet, artificial RARA homodimers inefficiently initiate APL, even in *Pml* null cells, which implies that PML fusion to RARA entails an unidentified gain of function central to leukemogenesis (Licht, 2006; Sternsdorf et al., 2006; Voisset et al., 2018).

Sumoylation is a reversible post-translational modification implicated in stress response. Sumoylation is mediated by UBC9, the universal SUMO E2 ligase, in an enzymatic cascade similar to ubiquitination (Celen and Sahin, 2020). In the context of chromatin, sumoylation is associated to transcriptional repression (Tharuka et al., 2025). PML is a massively sumoylated protein which serves as a scaffold to drive SUMO2-conjugation of a broad variety of nuclear body-associated partner proteins, particularly upon stress (Sahin et al., 2014; Tessier et al., 2022). SUMO2 conjugation was also linked to stress-induced target catabolism, as exemplified by ATO-induced PML/RARA degradation (Jaffray et al., 2023; Jaffray et al., 2025; Lallemand-Breitenbach et al., 2008; Lallemand-Breitenbach et al., 2001). Sumoylation often occurs on multiple members within key regulatory complexes, favoring their stability and regulating their functions (Psakhye and Jentsch, 2012). For example, the nuclear corepressor complex, assembled by NCoR1/2, may undergo SUMO-facilitated nuclear receptor association and/or transcriptional repression (Tiefenbach et al., 2006).

Here we demonstrate that fusion PML to RARA drives sumoylation of RARA-associated corepressors to promote transcriptional repression and progenitor immortalization.

## Results and discussion

### A key role for repression in hematologic progenitor immortalization by PML/RARA

In primary murine hematopoietic progenitors, PML/RARA expression results in their immortalization, as does over-expression of RARA, but not RARB or RARG (Fig. 1 A)(Du et al., 1999; Zhu et al., 2005). Yet, note that only PML/RARA ensures long-term progenitor immortalization after the firth replating, contrasting with RARA (Fig. 1A). Interestingly, at the second passage, the levels of RARA protein expression were consistently much higher than those of PML/RARA (Fig. 1 B), suggesting that the higher RARA expression levels in immortalized progenitors, compensate for its lower intrinsic repressive activity. Importantly, fusion of the POZ repression domains of PLZF onto RARB allowed efficient progenitor immortalization *ex vivo* (Fig. 1 A), highlighting the need of a strong basal transcriptional repression of RAR target genes in this process.

**Figure 1.**
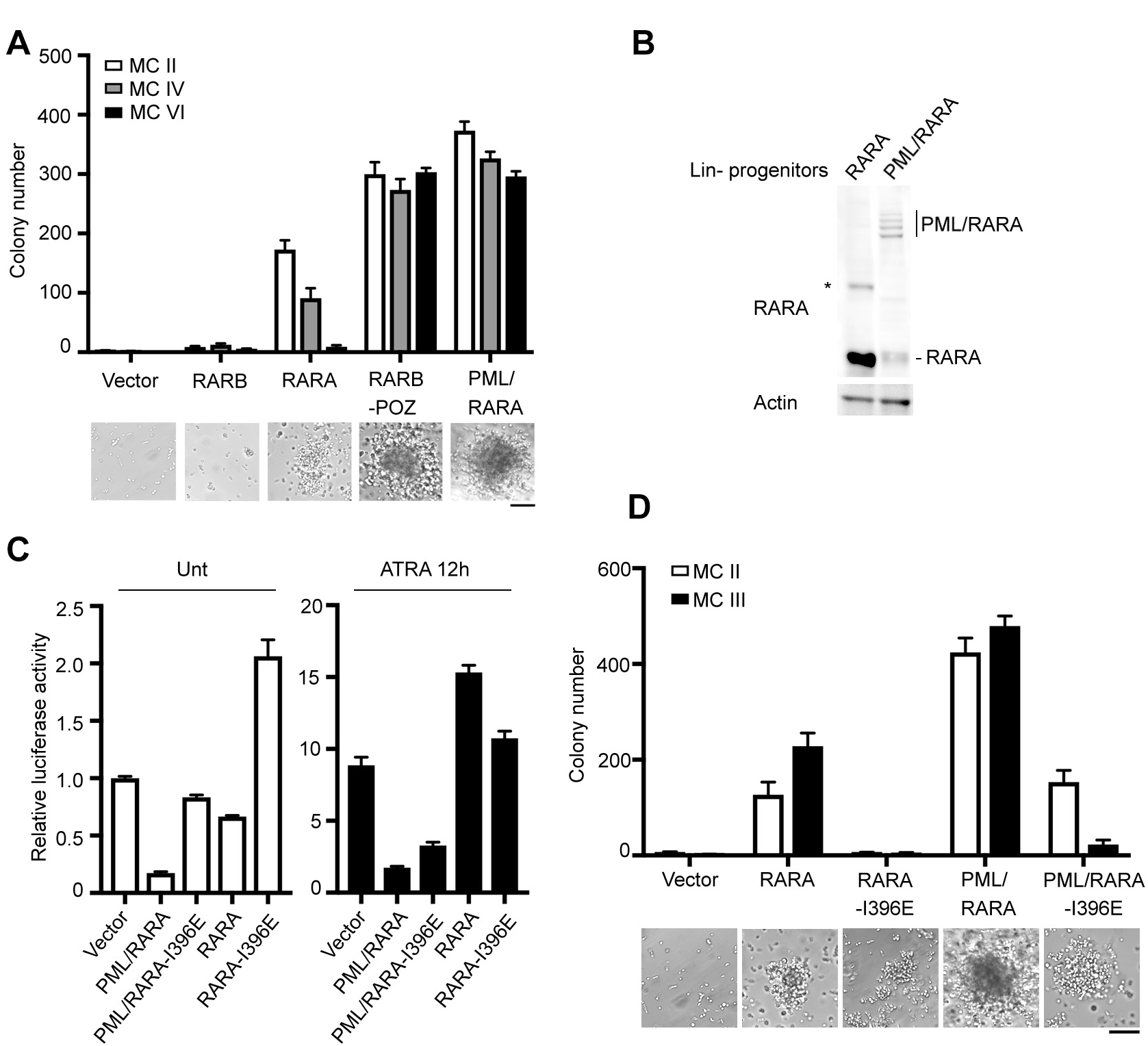
Transcriptional repression is critical for RARA and PML/RARA-driven progenitor immortalization. (**A**) Quantification of colony numbers and representative morphology (MC III) of primary mouse Lin^-^ hematopoietic progenitors transformed with RARA, RARB and indicated fusions. MC: serial plating in methyl-cellulose. Scale bar, 100 μm. (**B**) Western blot analysis of RARA and PML/RARA protein expression in transformed Lin^-^progenitors (MC II). (**C**) RARE/DR5 luciferase reporter assay from transfected 293T cells showing transcriptional activity of RARA and PML/RARA mutants after 12h all-trans-retinoic acid (RA) treatment. **(D)** Colony counts from serial replating assays and representative morphology (MC III) of primary mouse Lin^-^ hematopoietic progenitors transformed with RARA fusions or their I398E mutants that cannot bind NCoR. (**A**), (**C**) and (**D**), Data represent mean ± SD of triplicates. Representative of three independent experiments.

In RA absence, PML/RARA or RARA potently bind the NCOR1/2 complexes and RARA I396, is required for this binding (le Maire et al., 2010). Importantly, *PML/RARA^I396E^* or *RARA^I396E^* failed to efficiently repress basal expression of RA-sensitive reporters and no longer immortalized primary progenitors (Fig. 1, C and D). Collectively, repression of retinoic acid receptor target genes by NCoR-bound PML/RARA or RARA is required for primary hematopoietic progenitor immortalization, but can be mimicked by fusion of a strong repressive domain onto other retinoic acid receptors.

### Sumoylation of PML/RARA partners tightly correlates with transformation

To identify partners involved in enhanced repression by PML/RARA, we expressed a PML/RARA-BioID fusion in mouse Trp53^-/-^ HPC-7 stem/progenitor cells. Comparison between NLS-BioID and PML/RARA-BioID interactants identified 238 specific interactors with a high confidence (Fig. 2 A, Fig. S1 A; and Table S1). Fifty-five of them were very recently identified using a similar approach in different cellular system (Katerndahl et al., 2024)(Fig. S1 B). As expected, these comprised many known RARA or PML partners. The greatest enrichments were observed for RARA interactors with repressive abilities, including NCoR1/2, NRIP, GPS2, HDACs, SUMO2 (Fig. 2 A). RARA-associated proteins driving transcriptional activation (NCoA2/3) were also identified (Fig. 2 A). Unexpectedly, we also identified mRARA/B/G, which are not considered to be direct RARA interactors. PML binders (SUMO1/2, PIAS, TET2), which may modulate PML/RARA transcriptional output, were efficiently purified, together with some proteins (SKI, GPS2) proposed to be interactors of both PML and RARA. In biochemical validation experiments, RA indeed led to the expected dissociation of NCoR or GPS2, but greatly enhanced interactions with NCOA and NRIP1 (Fig. 2 B). A significant number of these PML/RARA interacting proteins were recently identified by SUMO2 proteomics in ATO-treated APL cells *in vivo*, notably NCoR1/2 and NRIP1 (Fig. 2 C and Fig. S1 C)(Tessier et al., 2022). Thus, similar to PML partners (Sahin et al., 2014), RARA interacting proteins may undergo ATO-enhanced SUMO2 conjugation in the presence of PML/RARA, most likely through PML-mediated tethering of UBC9. Accordingly, in transfected 293T cells, PML/RARA, but not RARA, allowed NCoR conjugation by SUMO2 (reversed by the SUMO inhibitor, TAK-981 (Lightcap et al., 2021))(Fig. 2 D and E), even in ATO absence. Interestingly, this was not the case for SUMO1.

**Figure 2.**
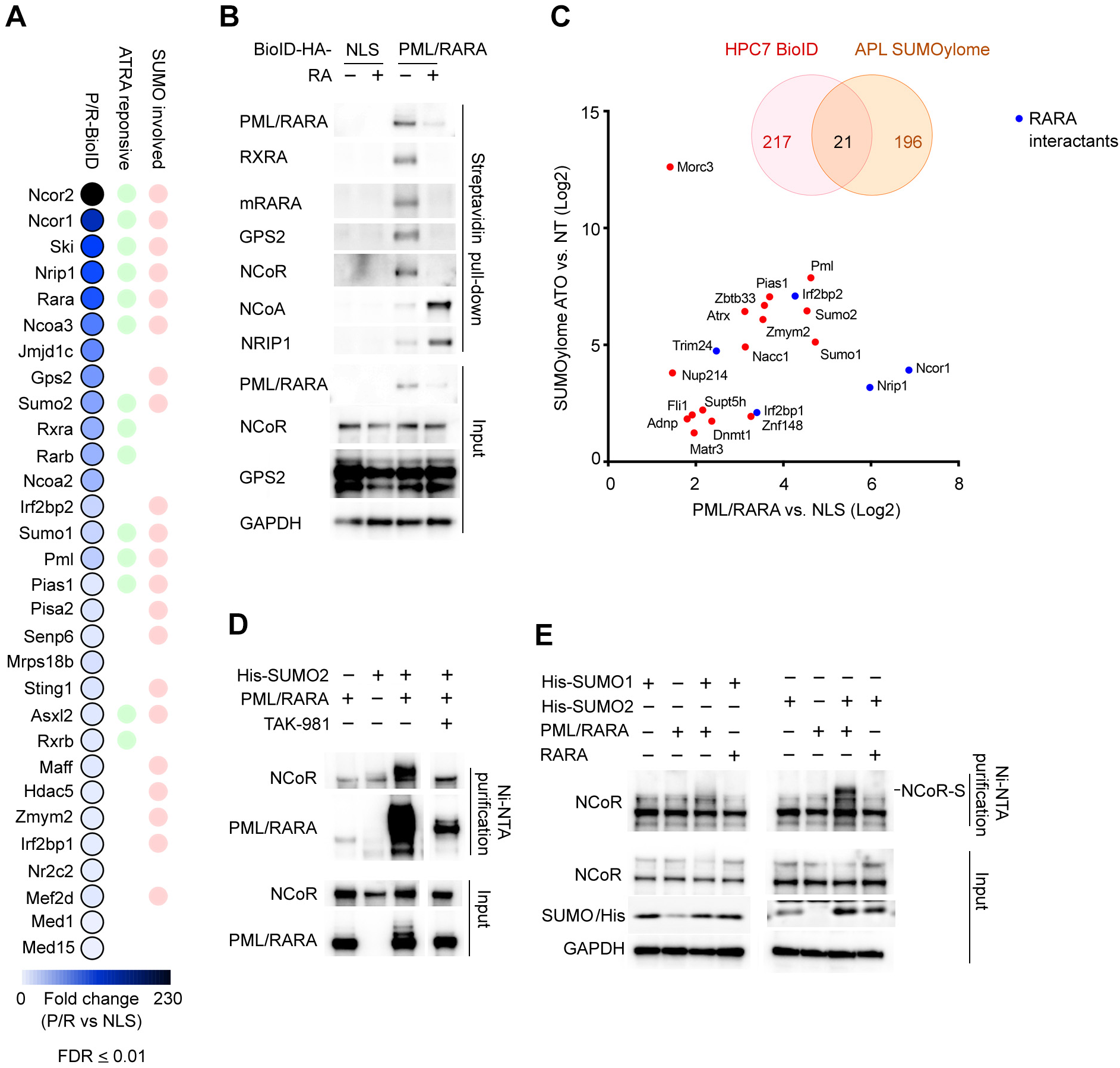
PML/RARA partners may undergo SUMO2 conjugation. (**A**) Dot plot of the top 30 proteins that showed increase of interaction with PML/RARA, ranked by fold enrichment over NLS-BioID control. N=2 for NLS-BioID, n=3 for PML/RARA-BioID. (**B**) Western blot validation of selected PML/RARA interactors after biotin labelling followed by streptavidin pulldown. Cells were treated with or without RA for 18h. (**C**) Dot plot of PML/RARA partner proteins previously found in arsenic (ATO)-responsive SUMOylome in primary mouse APL blasts. Known RARA binding protein are highlighted in blue. (**D**) and (**E**) Western blot analysis of lysates and Ni-NTA purified fractions from 293T cells transfected with indicated constructs. In (**D**), 293T cell were treated with SUMO inhibitor TAK-981 for 4h at day 2 after transfection before analysis.

To confirm that RARA interactants may undergo basal PML/RARA driven SUMO2 conjugation in stable transfectants, we then expressed PML/RARA-BioID and His_10_-SUMO2 in HPC7 cells and performed a stringent dual Streptavidin + Ni-NTA purification to identify SUMO2-conjugated PML/RARA interactants by mass spectrometry (Fig. 3 A). NCoR1/2 were again the top hits, together with other RARA-associated master regulators including RXRA, GPS2, GATA2, IRF2BP2, NSD2 and HDAC3/5/7/9 (Fig. 3B and C), several of which were known to undergo sumoylation (Fig 3D)(Andrade et al., 2016; Barysch et al., 2021; Chun et al., 2003), while other interactors were associated with PML nuclear bodies. To verify that PML/RARA may favor basal sumoylation of RARA-associated partners in a physiological setting, we used progenitors derived from His_10_-SUMO_3_ knock-in mice expressing, or not, PML/RARA at the preleukemic stage (where PML/RARA expression is barely detectable) or after evolution to full-blown APL (where PML/RARA becomes abundantly expressed (Westervelt et al., 2003)). These primary leukemic cells showed highly enhanced basal SUMO2-conjugation of PML/RARA-associated proteins, particularly NCoR and its associated proteins such as GPS2, as well as GATA2 or IRF2BP2 (Fig. 3 E), while sumoylation of Pml or Pml-associated proteins was decreased.

**Figure 3.**
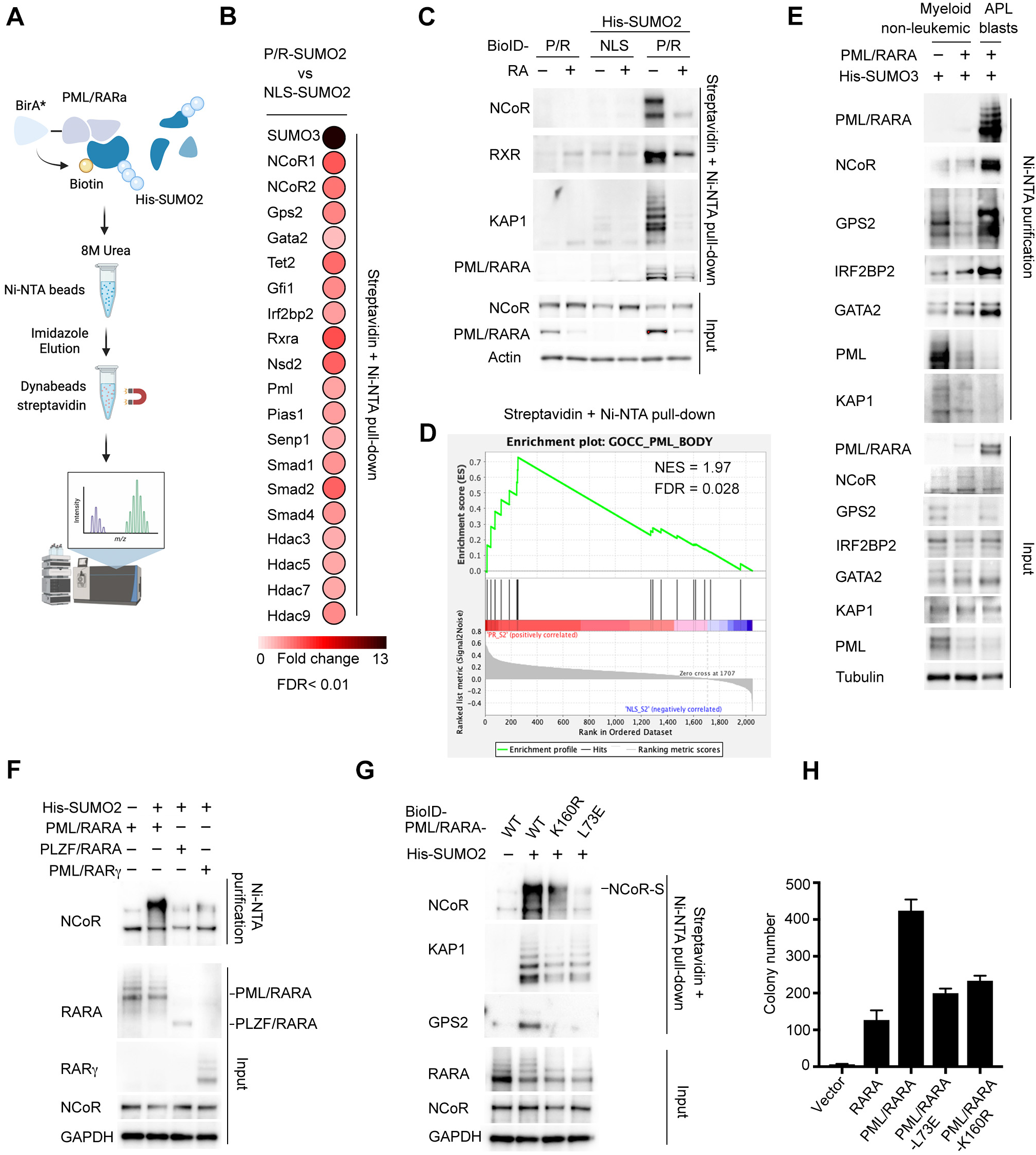
Fusion of PML to RARA allows RARA partners SUMOylation. (**A**) Experimental schematic of Ni-NTA + streptavidin double purification for mass spectrometry analysis from HPC7 cells expressing His_10_-SUMO_2_ and BioID proteins. (**B**) Dot plot of the top 20 PML/RARA interactors showing increased SUMO2 conjugation in His_10_-SUMO_2_-expressing HPC7 cells, comparing PML/RARA-BioID to NLS-BioID. n=4. (**C**) Western blot validation of lysates and Ni-NTA + Streptavidin purified fractions from HPC7 expressing PML/RARA-BioID alone, or with His_10_-SUMO_2_ plus NLS-BioID or PML/RARA-BioID. (**D**) GSEA analysis of proteomic data from His_10_-SUMO_2_ HPC7 cells expressing PML/RARA-BioID vs NLS-BioID. **E**) Western blot analysis of His_10_-SUMO_3_ knock-in mice expressing PML/RARA at non-leukemic stage or full-blown APL blasts. (**F**) Western blot analysis of lysates and Ni-NTA purified fractions from 293T cells transfected with His_10_-SUMO_2_ and indicated fusion constructs. (**G**) Western blot analysis of lysates and Ni-NTA + Streptavidin double purified fractions from 293T cells transfected with His_10_-SUMO_2_ and indicated PML/RARA mutants. (**H**) Colony counts of Lin^-^ progenitors transformed with RARA and PML/RARA mutants (MC III).

Among APL-associated fusion proteins, NCoR sumoylation was only observed upon PML/RARA-, but not PLZF/RARA- or PML/RARG- expression (Fig. 3 F), in keeping with the fact that PLZF does not bind UBC9, while RARG does not efficiently recruits NCoR (Farboud et al., 2003). Moreover, PML/RARA point mutants impaired in their UBC9 recruitment ability (L73E or K160R) (Bregnard et al., 2022; Wang et al., 2018; Zhu et al., 2005) yielded a significant decrease in SUMO2 conjugation of RARA-bound NCoR or GPS2 (Fig. 3 G). Importantly, these mutants were previously reported to be defective for PML/RARA-driven transformation (Fig. 3 H) (Wang et al., 2018; Zhu et al., 2005), tightly correlating efficient NCoR sumoylation to *PML/RARA*-driven transformation.

Collectively, the fusion of PML to RARA allows efficient SUMO2 conjugation of multiple RARA partners, notably the NCoR complex, potentially modulating its transcriptional output (Tiefenbach et al., 2006) and contributing to repression-mediated immortalization.

### Fusion of UBC9 to RARA allows long-term progenitor immortalization

If a major role of PML is to recruit UBC9 and enforce sumoylation of RARA-associated repressors, UBC9/RARA should be more efficient than RARA in its ability to immortalize progenitors. Indeed, UBC9/RARA fusions, but not overexpressed RARA or UBC9/RARA^I396E^, allowed long-term immortalization (Fig. 4 A). Remarkably, UBC9/RARA fusion boost self, NCoR, GATA2, IRF2BP2, GPS2 and RARA-partners sumoylation in both 293T transfectants and Lin^-^ progenitors stably expressing His_10_-SUMO2 (Fig. 4 B and C), and drives potent transcriptional repression (Fig. 4 D). Moreover, similar to PML/RARA, only very low levels of UBC9/RARA suffice to confer long-term replating ability (Fig. 4 E). Interestingly, UBC9/RARA^I396E^ still exerted some repressive activities and promoted some self-renewal, sharply contrasting with RARA^I396E^. Sumoylation of chromatin contributes cell identity and function (Theurillat et al., 2020). The UBC9/RARA fusion directly anchor the sumoylation machinery onto RARA binding sites and could exert broader control on gene expression, than solely through SUMO2 conjugation of NCoR or other RARA-associated proteins (Fig. 3B). Collectively, anchoring of UBC9 onto RAR target genes promotes their repression and drives progenitor immortalization.

**Figure 4.**
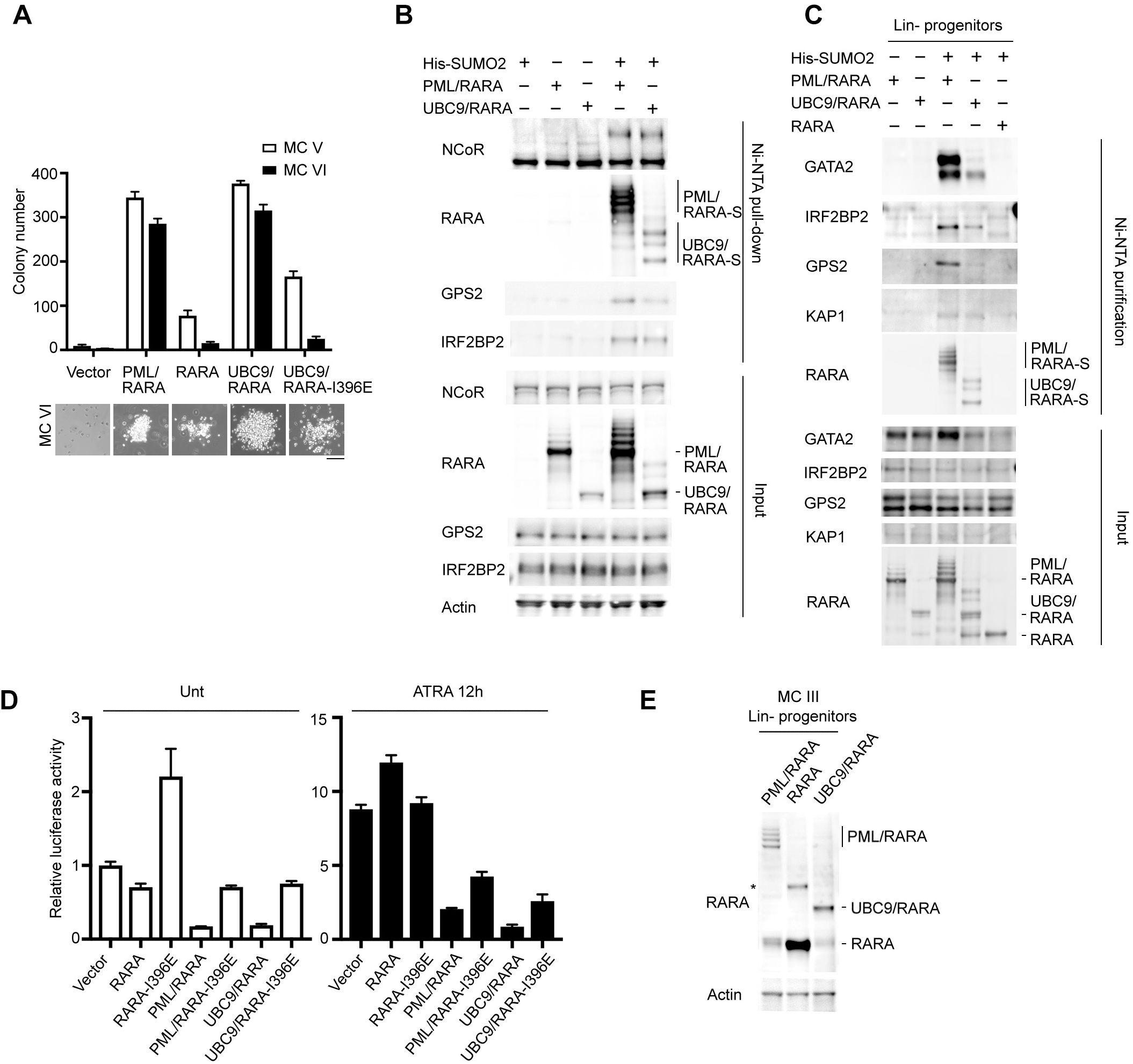
Fusion of UBC9 to RARA enforces sumoylation of RARA-bound partners and allows long-term progenitors immortalization. (**A**) Colony counts from serial replating assays and representative morphology (MC VI) of primary mouse Lin^-^ hematopoietic progenitors transformed with RARA fusions or their I398E mutants defective for NCoR binding. (**B** and **C**) Western blot analysis of lysates and Ni-NTA purified fractions from 293T cells transfected with indicated constructs (**B**) or progenitors transformed with RARA fusions (**C**). (**D**) RARE/DR5 luciferase reporter assay showing transcriptional activity of RARA and PML/RARA mutants after 12h RA treatment. (**E**) Western blot analysis of RARA, PML/RARA and UBC9/RARA protein expression in transformed Lin^-^ progenitors.

### Sumoylation inhibitors drive target gene activation, differentiation and loss of self-renewal

If NCoR sumoylation has any significant role in PML/RARA-dependent transcriptional repression, chemical inhibitors of sumoylation (Lightcap et al., 2021) should activate PML/RARA or UBC9/RARA targets. TAK-981 treatment indeed yielded re-activation of canonical RAR targets (Fig. 5 A and Fig. S2 A), in PML/RARA or UBC9-RARA-transformed primary progenitors, but not in RARA- or PLZF/RARA-immortalized ones. As expected, TAK-981 activated IFNγ genes in both settings (Fig. S2 B). Thus, repression driven by these fusions directly involves sumoylation of NCoR, and other sumoylated PML/RARA interactors. Moreover, *ex vivo t*reatment of PML/RARA or UBC9-RARA-transformed primary progenitors by TAK-981 induced their differentiation and stemness lost (Fig. 5 B, C and D). TAK-981-initiated differentiation also was obtained *in vivo*, using murine APL models (Fig. 5 E) and was accompanied by target gene re-activation (Fig. 5 F).

**Figure 5.**
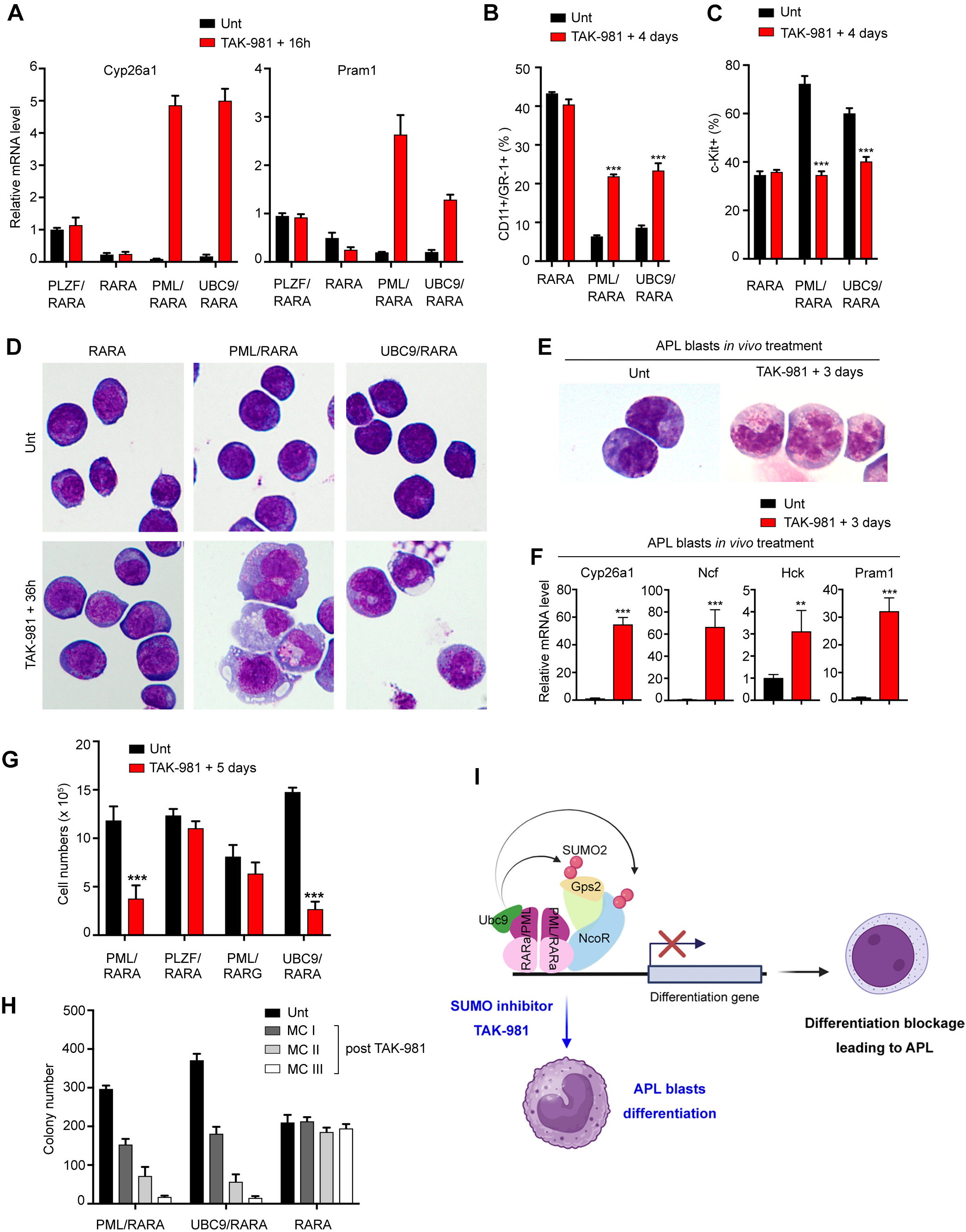
Inhibition of SUMO conjugation re-activates PML/RARA target genes and induces differentiation. (**A**) Real-time PCR for RARA targets, *Cyp26a1* and *Pram1*, gene expression in primary progenitors transformed with indicated constructs post TAK-981 (100 nM) treatment for 18h. (**B** and **C**) FACS analysis of differentiation (**B**) or stemness marker (**C**) expression in progenitors transformed with indicated fusions after 4 days TAK-981 (50 nM) treatment. (**D**) MGG staining of progenitors as (**B** and **C**) after 36h TAK-981 treatment. (**E**) MGG staining of primary mouse APL blasts treated with TAK-981 *in vivo* for 3 days. (**F**) Real-time PCR for RARA targets, *Cyp26a1*, *Pram1, Ncf,* and *Hck* gene expression in APL blasts treated with TAK-981 as in (**E**). (**G**) Abundance of Lin^-^ progenitors expressing the indicated fusions after 5 days of TAK-981 (50 nM) treatment. (**H**) Colony counts of primary progenitors transformed with indicated fusions. Cells were treated with TAK-981 (50 nM) for 5 days then replating subsequently in methylcellulose without SUMO inhibitor. (**I**) Model scheme illustrating PML-driven sumoylation of PML/RARA-bond repressors driving the differentiation block in APL. (**A**, **B, C, F, G** and **H**) Data represent mean ± SD of triplicates; unpaired t test: *p < 0.05, **p < 0.005, ***p < 0.001; representative of three independent experiments.

We then explored the long-term effects of TAK-981 on clonogenic activity of progenitors transformed with various RARA-fusions. A 5 days *ex vivo* treatment drove a significant decrease in the number of clones (Fig. S2 C) and an even sharper reduction in total number of cells, solely in PML/RARA- and UBC9/RARA-transformed cells (Fig. 5 G). Remarkably, subsequent replating of these TAK-981-treated progenitors led to a progressive exhaustion of their clonogenic activities, while RARA-transformed progenitors were unaffected (Fig. 5 H). Collectively, inhibition of sumoylation in PML- or UBC9- RARA fusions reactivates RAR signaling to drive APL growth arrest and differentiation.

## Discussion

Retinoid signaling remains an intense field of investigations (Esposito et al., 2024). Transcriptional repression of retinoic acid target genes was proposed to underlie APL pathogenesis (de The et al., 2017), as formally demonstrated here by the *PML/RARA^I396E^* mutant, using progenitor immortalization as a surrogate for transformation. In that respect, in PLZF/RARA variants APLs, the PLZF POZ domain yields a super-repressive phenotype that entails clinical RA-resistance (Licht et al., 1995). Why PML is the recurrent fusion partner and how this relates to transcriptional repression remained poorly understood (Sternsdorf et al., 2006). We identify a novel mechanism for PML/RARA-driven hyper-repression, through PML-mediated sumoylation of PML/RARA-bound proteins, particularly the NCoR complex, whose stability, nuclear receptor binding and repressive ability are all tightly regulated by sumoylation (Fig. 5 I) (Hua et al., 2016; Tiefenbach et al., 2006). Importantly, we also identified other sumoylated key RARA-associated regulatory proteins such GATA2, IRF2BP2, NSD2… (Fig. 3B) whose activities are regulated by sumoylation (Andrade et al., 2016; Barysch et al., 2021; Chun et al., 2003), suggesting that their PML-driven sumoylation may also impinge on PML/RARA function. IRF2BP2 may be a RARA fusion partner in very rare APLs (Shimomura et al., 2016; Yin et al., 2015), while GATA2 forms tight complexes with RARA on DNA (Katerndahl et al., 2024; Tsuzuki et al., 2004). More broadly, further chromatin sumoylation around PML/RARA binding sites may entail transcriptional repression (Stielow et al., 2008).

Supporting the relevance of sumoylation to PML/RARA-mediated repression and function, domains of PML involved in UBC9 recruitment and sumoylation control (RING, K160) were also required for enhanced NCoR sumoylation, target gene regulation, TAK-981 sensitivity and, critically, leukemogenesis *in vivo* (Wang et al., 2018; Zhu et al., 2005). NCoR complex sumo-interacting motifs may also boost its own binding onto sumoylated PML/RARA, as suggested for some other nuclear receptors (Hua et al., 2016; Paakinaho et al., 2021), further contributing to the enhanced association of this master repressive complex and the fusion protein. That UBC9/RARA exerts much stronger transforming effects than RARA and confers phenotypic sensitivity to UBC9 inhibitors strongly argues for a direct functional role of sumoylation of RARA partners in transcriptional repression, independently from any specific PML interactions, for example with the Daxx repressor. Finally, in addition to sumoylated RARA partners (NCoR1/2, GPS2, GATA2, Irf2bp2, Nsd2…), we also identified other interactors with potential importance in APL pathogenesis, such as the Ski oncogene, which binds to RARA to inhibit RA signaling (Ritter et al., 2006) and confers RA-sensitivity to immortalized progenitors (Dahl et al., 1998; Melling et al., 2013).

PML fusion to RARA drives dimer-dependent gains of function, including relaxed DNA-binding specificity, enhanced corepressor association (Kamashev et al., 2000; Sternsdorf et al., 2006) or recruitment of PML-bound partners (Occhionorelli et al., 2011; Zhu et al., 2005). Previous studies suggested that these are insufficient to drive immortalization on their own (Sternsdorf et al., 2006; Voisset et al., 2018). The experiments reported here illustrate how critical PML/UBC9-driven post-translational modifications within the oncogenic complex can contribute to transformation, a key gain of function resulting from the fusion of the PML and RARA genes. Such cross-talks within the fusion to favor target gene repression explain with PML and RARA are the preferred targets of translocations driving APL. Interestingly, sumoylation was recently shown to modulate oncogenesis in other settings (Li et al., 2025; Zhang et al., 2025), but also to favor therapeutic response through nuclear receptors activation (Valima et al., 2025), opening novel perspectives for sumoylation inhibitors in cancer therapy, beyond boosting of immunotherapies (Lightcap et al., 2021).

## Methods

### Cell Lines

Mouse HPC7 Trp53^-/-^ stem/progenitor cell line used in this study was generated from the parental HPC7 cell line (a kind gift of Camille Lobry) by using the Alt-R CRISPR/Cas9 technology (IDT). Briefly, HPC7 cells were electroporated with guide RNA targeting TP53 (5’GCGCTGACCCACAACTGCAC) and recombinant Cas9 reagent following the recommendation of the manufacturer. Single-cell cloning was performed into 96 wells and knockout of the Trp53 gene was confirmed by sequencing, immunofluorescence and Western blotting analysis (data not shown). Trp53^-/-^ HPC7 cells were retrovirally transduced with PML/RARA-/NLS- HA-BioiD and/or not His_10_-SUMO2 vectors. All HPC7 cell lines were maintained in Iscove’s modified Dulbecco’s medium plus GlutaMax (Cat # 31980030; Thermo Fisher Scientific) supplemented with 5% FBS in the presence of 100 U/mL of penicillin, 100 μg/mL of streptomycin, 0.1 mM 2-mercaptoethanol and 100ng/ml of soluble Kit ligand recovered from CHO cell line stably expressing mouse c-Kit. HEK293T cells were maintained in Dulbecco’s Modified Eagle Medium plus GlutaMAX (Cat # 41965062; Thermo Fisher Scientific) supplemented with 10% FBS, 100 U/mL of penicillin and 100 μg/mL of streptomycin. Exploration of retinoic acid response using the RARE/DR5 was performed as previously (de Thé et al., 1990).

### Plasmid constructs

For pMSCV-Flag-HA-POZ-RARB plasmid, nested PCR was used to amplify a Flag-HA tag in frame with POZ domain from MSCV-PLZF-RARA with the following primers: Flag-HA-POZ For:5’GCTGAATTCGCCACC*ATGG*ACTACAAGGACGACGATGACAAGCTCGATGGAGGATAC CCCTACGACGTGCCCGACTACGCCGATCTGACAAAAATGGGCATGATCCAGCTGCAGAACCCTAGCCAC 3’ ; POZ rev: 5’ GGC<u>TCTAGA</u>CCCGCCTCCACCGATGGTCTCCAGCATCTTC 3’. The Flag-HA-POZ PCR product including a glycine-serine linker was then cloned in-frame with RARB into pMSCV-RARB retroviral vector (a kind gift of Albane le Maire).

To construct UBC9-RARA expression plasmids, UBC9 was PCR amplified from pSG5-Ubc9 plasmid and cloned in-frame with RARA in both pMSCV-RARA and pCMV-RARA vectors using the following primers: UBC9-MSCV-For TCTCTCGAGGCCACCATGTCGGGGATCGC; UBC9-MSCV-Rev TTCGTTAACTCACGGGGAGTGGGTGGC; CMV-UBC9-For CTGGCTAGCC ACCATGTCGGGGATCG; CMV-UBC9-Rev ACCGGATCCCGGGGAGTGG GTGGCC

### Colony formation assay

For methylcellulose colony formation assay, primary murine Lin^-^ progenitors were obtained from 5-fluorouracil-treated C57Bl/6Nj mice using lineage cell depletion kit (cat# 130-090-858; Miltenyi Biotec). Mice were purchased from Janvier Labs. Lin^-^ progenitors were retrovirally transduced with indicated fusions and maintained in methylcellulose (Cat # 3231; Stemcell Technology) supplemented with 10 ng/ml of IL-3, IL-6, GM-CSF, and 50 ng/ml of SCF as previous described (Zhu et al., 2005), and selected by puromycin (1 μg/ml) or neomycin (500 μg/ml). To determine the impacts of fusion proteins and TAK-981 (HY-111789; MedChemExpress) on progenitors’ immortalization, 5,000 cells per well in 6-cell plate were cultured and subsequently re-plated to assess their clonogenic capacity.

### Biochemistry

To purify PML/RARA interactants or His_10_-SUMO1/2 conjugated proteins, HPC7, Lin^-^progenitors, and 293T cells were transduced/transfected with indicated plasmids and lysed by buffer A containing 8 M urea, 50 mM Tris pH 8, 150 mM NaCl, 10 mM NEM (cat # E3876; Sigma), 15 mM imidazole, universal nuclease (cat # 88702; Thermo Scientific) and proteinase inhibitors (cat # 11 836 170 001; Roche). Protein concentration was determined using Bradford reagent (cat # B6916; Sigma). Equal amount of protein lysates containing 40 uM PR-619 (cat # 662147; Sigma) were incubated with nickel-nitrilotriacetic (Ni-NTA) agarose beads (cat # L30210; Qiagen) or streptavidin Dynabeads (Cat # 65602; Thermo Fisher Scientific) overnight at 4°C. The beads were washed six times with buffer A and then analyzed by Western blot.

For streptavidin + Ni-NTA double-purification experiments, His_10_-SUMO_2_ conjugated proteins were eluted by RIPA buffer containing 250 mM imidazole, 50 mM Tris (pH 8.0), 0.15 M NaCl, 1% NP40, 1% sodium deoxycholate, 0.1% SDS, proteinase inhibitor, and 40 μM PR-619. Biotinylated and SUMO2ylated protein were enriched by streptavidin Dynabeads overnight at 4°C. The beads were washed three times with RIPA buffer and then analyzed by Western blot.

The following antibodies were used with 1:1000 dilution for immunodetection in precipitation samples and cell lysates: Actin (A2066; Sigma); RARA and PML (home-made affinity-purified rabbit and mouse); GPS2 and NCoR (PA5-76547, A301-145A; Thermo Fisher Scientific); NRIP1 (NBP3-12264; Novus); NCOA, RXRA, tubulin, GATA2, SUMO1, and SUMO2 (ab10491, ab125001, ab4074, ab109241, ab32058, ab81371; Abcam); KAP1 and His tag (4124, 2365; Cell signaling); IRF2BP2 (18847-1-AP; Proteintech); TET2 (GTX124205; GeneTax).

### RNA analyses

Total RNA extraction from progenitors and APL blasts were performed with the RNeasy Plus Mini Kit (cat# 74134; Qiagen) and quantified using a Nondrop One-One. cDNA was prepared from 1 ug of total RNA with an iScript^TM^ cDNA Synthesis kit (cat#1708891; Bio-Rad). Quantitative real-time PCR on cDNA was performed on Bio-Rad CFX thermal cycles with TaqMan Gene Expression assays from Life Technologies (*Gapdh*, 4352339E; *Cyp26a1*, Mm00514486_m1; *Pram1*, Mm02744730_g1; *Ncf1*, Mm00447921_m1; *Hck*, Mm01241463_m1; *Ifnγ,* Mm01168134_m1)

### Flow cytometry analysis

Progenitor population were analyzed using FACSCANTO II (BD). Single-cell suspensions were blocked with Fc-Block (BD) for 15 min on ice. The following antibodies were used to stain fusion protein transformed Lin^-^ progenitor cells (1:1000): APC-conjugated CD117 (Clone 2B8; eBioscience), APC-Cy7 conjugated anti-CD11b (Clone M1/70, BioLegend), and phycoerythrin-conjugated anti-Gr1 (Clone RB6-8C5, eBioscience). Staining was performed overnight at 4°C. Cells were washed and resuspended in phosphate buffer saline (PBS) with paraformaldehyde 0.2% before FACS analysis.

### Mouse model

His_10_-HA-SUMO_3_ Knock-in strain was generated using the CRISPR/Cas9 system by the i-GONAD electroporation method with the BALB/cByJ strain. BALB/cByJ mice were purchased from Charles River. crRNA, tracrRNA, ssDNA, and Cas9 nuclease were purchased from IDT (Crispr target sequence: TCAGTTTCCATTTCTTGAT; ssODN sequence: CATTTCCCGCCTTCACAGACCTAATAAGAAATGCATCACCA TCACCATCATCATCATCATC ATTACCCATACGATGTTCCAGATTACGCTGAAACTG AACCAGTTTCCGTGCAGAAGGTAC CTGCACCC). To obtain His_10_-HA-Sumo_3_ *Pml/Rara* double mutant mice, this strain was crossed with Ctsg-PML/RARA mice on a BALB/cByJ background. For Fig 3D, APL blasts were collected from the bone marrow. Non-leukemic myeloid cells were enriched by depletion of lymphoid lineage using CD4/CD8, CD49b, and CD19 microbeads (cat# 130-116-480, 130-052-501, 130-121-301) and MACS column from Miltenyi Biotec.

Leukemic blasts in Fig 5E and F were derived from h-MRP8-*PML/RARA* transgenic mice and transplanted by intravenous injection into FVB mice. Serial intravenous transplantations were performed using 10^4^ GFP sorted APL cells from bone marrow. Animals were handled according to the guidelines of institutional animal care committees using protocols approved by the “Comité d’Ethique Experimentation Animal Paris-Nord (no.121). Mice were maintained in a 12h light-dark cycle animal facility under specific pathogen-free conditions with free access to water and food (A03: SAFE; Institut de Recherche Saint Louis, Paris, France).

### Mass spectrometry analyses

Experiments were performed at the 3P5 proteomics facility of the Sorbonne University. Ni-NTA or Streptavidin purified proteins were gel purified and trypsin-digested overnight and processed essentially as described (Hospital et al., 2018). Mass spectrometric data acquisition was performed on a NanoElute II UHPLC system coupled to a TIMS-TOF-HT mass spectrometer (Bruker Daltonics), with a nano-electrospray ion source (Bruker CaptiveSpray source). All samples were acquired in a DIA-PASEF acquisition mode, with an m/z range 100 -1700 and an ion mobility (IM) range of 0,7 – 1,3 1/K0. Isolation windows were fixed at 35 Da, with a cycle time estimate of 1.17 s. Mass and IM calibration were done linearly using three ions from the Agilent ESI LC/MS tuning mix (m/z 622.0290, IM 0.9917 1/K0; m/z 922.0098, IM 1.1984 1/K0; m/z 1221.9906, IM 1.3934 1/K0. Mass spectrometry data was analyzed by the software DIA-NN (Data-Independent Acquisition by Neural Networks), version 1.8.1. Data filtering, statistical, and differential analyses were all carried out with the PROSTAR software, version 1.30.7. Data was refined by excluding missing lines and contaminants from the dataset, followed by log2 transformation of LFQ intensities. Normalization was carried out by the method of quantile centring, within each condition. Significant differences were evaluated by the Limma test, using an adaptive Benjamini-Hochberg correction. Proteins were deemed significant if p-values were consistent with an FDR cut-off of 1%.

### Bioinformatic and statistical analyses

Gene set enrichment analysis was performed using Gene ontology (C5.all) containing 16107 gene sets. PML and RARA interactome shown in Extended Data Fig. 1c was analysis with STRING and generated by Cytoscape (verson3.10.2). Dot plot shown in Fig.1e and 2a were generated by ProHits-viz. Graphpad Prism software was used to perform multiple unpaired two-tailed t-test to determine the *P* value and look for significant changes in the data generated from the fusions expressing cells and TAK-981 treatment *in vivo* or *ex vivo*. All data are expressed as mean ± s.d. For all graphs, \**P* = 0.01-0.05, \*\**P* = 0.001-0.01, and \*\*\**P* < 0.001.

**Supplementary materials** include a list of 238 significant PML/RARA associated protein identified in HPC7 cell lines (Sup. Table 1), data on PML/RARA-interacting proteins (Sup. Figure 1) and data on Tak-981 response in primary progenitors (Sup. Figure 2).

## Acknowledgements

Work in the laboratory is supported by the ERC (PML-Therapy, RARA-AML), as well as Institut National du Cancer (PLBio). The animal facility, was supported in part by ANR, through the France 2030 program, ANR-23-IAHU-0005, Paris St. Louis Leukemia Institute. We warmly thank T. Ley (Washington University, St. Louis) for his generous gift of CatG-*PML::RARA^+/+^* C57BL/6 mice (Westervelt et al., 2003). We warmly thank the P3S proteomic facility of Sorbonne University for expert analyses and advise.

## Authors contributions

HCW, EL, CE, MNK, YZ and MCG performed experiments. HT, MCG and HCW wrote the paper which was reviewed and accepted by all co-authors.

## Competing interests

The authors declare no conflict of interest related to this work.

## Notes

### Competing Interest Statement

The authors have declared no competing interest.

## Reference

Andrade, D., M. Velinder, J. Singer, L. Maese, D. Bareyan, H. Nguyen, M.B. Chandrasekharan, H. Lucente, D. McClellan, D. Jones, S. Sharma, F. Liu, and M.E. Engel. 2016. SUMOylation Regulates Growth Factor Independence 1 in Transcriptional Control and Hematopoiesis. Mol Cell Biol 36:1438–1450.

Barysch, S.V., N. Stankovic-Valentin, T. Miedema, S. Karaca, J. Doppel, T. Nait Achour, A. Vasudeva, L. Wolf, C. Sticht, H. Urlaub, and F. Melchior. 2021. Transient deSUMOylation of IRF2BP proteins controls early transcription in EGFR signaling. EMBO Rep 22:e49651.

Bregnard, T., A. Ahmed, I.V. Semenova, S.K. Weller, and I. Bezsonova. 2022. The B-box1 domain of PML mediates SUMO E2-E3 complex formation through an atypical interaction with UBC9. Biophys Chem 287:106827.

Celen, A.B., and U. Sahin. 2020. Sumoylation on its 25th anniversary: mechanisms, pathology, and emerging concepts. FEBS J 287:3110–3140.

Chun, T.H., H. Itoh, L. Subramanian, J.A. Iniguez-Lluhi, and K. Nakao. 2003. Modification of GATA-2 transcriptional activity in endothelial cells by the SUMO E3 ligase PIASy. Circ Res 92:1201–1208.

Dahl, R., M. Kieslinger, H. Beug, and M.J. Hayman. 1998. Transformation of hematopoietic cells by the Ski oncoprotein involves repression of retinoic acid receptor signaling. Proc. Natl. Acad. Sci. USA 95:11187–11192.

de The, H., C. Chomienne, M. Lanotte, L. Degos, and A. Dejean. 1990. The t(15;17) translocation of acute promyelocytic leukaemia fuses the retinoic acid receptor alpha gene to a novel transcribed locus. Nature 347:558–561.

de Thé, H., C. Lavau, A. Marchio, C. Chomienne, L. Degos, and A. Dejean. 1991. The PML-RAR alpha fusion mRNA generated by the t(15;17) translocation in acute promyelocytic leukemia encodes a functionally altered RAR. Cell 66:675–684.

de The, H., P.P. Pandolfi, and Z. Chen. 2017. Acute Promyelocytic Leukemia: A Paradigm for Oncoprotein-Targeted Cure. Cancer Cell 32:552–560.

de Thé, H., M.d.M. Vivanco-Ruiz, P. Tiollais, H. Stunnenberg, and A. Dejean. 1990. Identification of a retinoic acid responsive element in the retinoic acid receptor beta gene. Nature 343:177–180.

Dos Santos, G.A., L. Kats, and P.P. Pandolfi. 2013. Synergy against PML-RARa: targeting transcription, proteolysis, differentiation, and self-renewal in acute promyelocytic leukemia. The Journal of experimental medicine 210:2793–2802.

Du, C., R.L. Redner, M.P. Cooke, and C. Lavau. 1999. Overexpression of wild-type retinoic acid receptor alpha (RAR alpha) recapitulates retinoic acid-sensitive transformation of primary myeloid progenitors by acute promyelocytic leukemia RAR alpha-fusion genes. Blood 94:793–802.

Esposito, M., J.K. Amory, and Y. Kang. 2024. The pathogenic role of retinoid nuclear receptor signaling in cancer and metabolic syndromes. J Exp Med 221:

Farboud, B., H. Hauksdottir, Y. Wu, and M.L. Privalsky. 2003. Isotype-restricted corepressor recruitment: a constitutively closed helix 12 conformation in retinoic acid receptors beta and gamma interferes with corepressor recruitment and prevents transcriptional repression. Mol Cell Biol 23:2844–2858.

Geoffroy, M.C., c. Esnault, and H. de The. 2021. Retinoids in haematology, a timely revival ?. Blood 137:2429–2437.

Hospital, M.A., A. Jacquel, F. Mazed, E. Saland, C. Larrue, J. Mondesir, R. Birsen, A.S. Green, M. Lambert, P. Sujobert, E.F. Gautier, V. Salnot, M. Le Gall, J. Decroocq, L. Poulain, N. Jacque, M. Fontenay, O. Kosmider, C. Recher, P. Auberger, P. Mayeux, D. Bouscary, J.E. Sarry, and J. Tamburini. 2018. RSK2 is a new Pim2 target with pro-survival functions in FLT3-ITD-positive acute myeloid leukemia. Leukemia 32:597–605.

Hua, G., L. Paulen, and P. Chambon. 2016. GR SUMOylation and formation of an SUMO-SMRT/NCoR1-HDAC3 repressing complex is mandatory for GC-induced IR nGRE-mediated transrepression. Proc Natl Acad Sci U S A 113:E626–634.

Jaffray, E.G., M.H. Tatham, B. Mojsa, M. Liczmanska, A. Rojas-Fernandez, Y. Yin, G. Ball, and R.T. Hay. 2023. The p97/VCP segregase is essential for arsenic-induced degradation of PML and PML-RARA. J Cell Biol 222:

Jaffray, E.G., M.H. Tatham, B. Mojsa, A. Plechanovova, A. Rojas-Fernandez, J.C.Y. Liu, N. Mailand, A.F.M. Ibrahim, G. Ball, I.M. Porter, and R.T. Hay. 2025. PML mutants from arsenic-resistant patients reveal SUMO1-TOPORS and SUMO2/3-RNF4 degradation pathways. J Cell Biol 224:

Kamashev, D.E., A.V. Balandina, and V.L. Karpov. 2000. Tramtrack protein-DNA interactions. A cross-linking study. J. Biol. Chem. 275:36056–36061.

Katerndahl, C.D.S., O.R.S. Rogers, R.B. Day, Z. Xu, N.M. Helton, S.M. Ramakrishnan, C.A. Miller, and T.J. Ley. 2024. PML::RARA and GATA2 proteins interact via DNA templates to induce aberrant self-renewal in mouse and human hematopoietic cells. Proc Natl Acad Sci U S A 121:e2317690121.

Lallemand-Breitenbach, V., M. Jeanne, S. Benhenda, R. Nasr, M. Lei, L. Peres, J. Zhou, J. Zhu, B. Raught, and H. de The. 2008. Arsenic degrades PML or PML-RARalpha through a SUMO-triggered RNF4/ubiquitin-mediated pathway. Nat Cell Biol 10:547–555.

Lallemand-Breitenbach, V., J. Zhu, F. Puvion, M. Koken, N. Honore, A. Doubeikovsky, E. Duprez, P.P. Pandolfi, E. Puvion, P. Freemont, and H. de The. 2001. Role of Promyelocytic Leukemia (PML) Sumolation in Nuclear Body Formation, 11S Proteasome Recruitment, and As(2)O(3)-induced PML or PML/Retinoic Acid Receptor alpha Degradation. J Exp Med 193:1361–1372.

le Maire, A., C. Teyssier, C. Erb, M. Grimaldi, S. Alvarez, A.R. de Lera, P. Balaguer, H. Gronemeyer, C.A. Royer, P. Germain, and W. Bourguet. 2010. A unique secondary-structure switch controls constitutive gene repression by retinoic acid receptor. Nat Struct Mol Biol 17:801–807.

Li, Z., K.E. Koch, D.T. Thompson, D.M. Van der Heide, J. Chang, C.M. Franke, M.O. Suraju, A.C. Beck, A.W. Lorenzen, J.R. White, N.I. Bartschat, M.V. Kulak, D.K. Meyerholz, C. Kenny, and R.J. Weigel. 2025. Sumoylated Etv1 establishes mouse mammary cancer stem cells that support tumorigenesis by non-stem cancer cells. Dev Cell 60:2264–2278 e2267.

Licht, J.D. 2006. Reconstructing a disease: What essential features of the retinoic acid receptor fusion oncoproteins generate acute promyelocytic leukemia? Cancer Cell 9:73–74.

Licht, J.D., C. Chomienne, A. Goy, A. Chen, A.A. Scott, D.R. Head, J.L. Michaux, Y. Wu, A. DeBlasio, W.H. Miller, Jr., and, et al. 1995. Clinical and molecular characterization of a rare syndrome of acute promyelocytic leukemia associated with translocation (11;17). Blood 85:1083–1094.

Lightcap, E.S., P. Yu, S. Grossman, K. Song, M. Khattar, K. Xega, X. He, J.M. Gavin, H. Imaichi, J.J. Garnsey, E. Koenig, H. Zhang, Z. Lu, P. Shah, Y. Fu, M.A. Milhollen, B.A. Hatton, J. Riceberg, V. Shinde, C. Li, J. Minissale, X. Yang, D. England, R.A. Klinghoffer, S. Langston, K. Galvin, G. Shapiro, S.M. Pulukuri, S.Y. Fuchs, and D. Huszar. 2021. A small-molecule SUMOylation inhibitor activates antitumor immune responses and potentiates immune therapies in preclinical models. Sci Transl Med 13:eaba7791.

Lin, R.J., and R.M. Evans. 2000. Acquisition of oncogenic potential by RAR chimeras in acute promyelocytic leukemia through formation of homodimers. Mol Cell 5:821–830.

Melling, M.A., C.R. Friendship, T.G. Shepherd, and T.A. Drysdale. 2013. Expression of Ski can act as a negative feedback mechanism on retinoic acid signaling. Dev Dyn 242:604–613.

Occhionorelli, M., F. Santoro, I. Pallavicini, A. Gruszka, S. Moretti, D. Bossi, A. Viale, D. Shing, S. Ronzoni, I. Muradore, M. Soncini, G. Pruneri, P. Rafaniello, G. Viale, P.G. Pelicci, and S. Minucci. 2011. The self-association coiled-coil domain of PML is sufficient for the oncogenic conversion of the retinoic acid receptor (RAR) alpha. Leukemia : official journal of the Leukemia Society of America, Leukemia Research Fund, U.K 25:814–820.

Paakinaho, V., J.K. Lempiainen, G. Sigismondo, E.A. Niskanen, M. Malinen, T. Jaaskelainen, M. Varjosalo, J. Krijgsveld, and J.J. Palvimo. 2021. SUMOylation regulates the protein network and chromatin accessibility at glucocorticoid receptor-binding sites. Nucleic Acids Res 49:1951–1971.

Psakhye, I., and S. Jentsch. 2012. Protein group modification and synergy in the SUMO pathway as exemplified in DNA repair. Cell 151:807–820.

Rerolle, D., H.C. Wu, and H. de The. 2024. Acute Promyelocytic Leukemia, Retinoic Acid, and Arsenic: A Tale of Dualities. Cold Spring Harb Perspect Med 14:

Ritter, M., D. Kattmann, S. Teichler, O. Hartmann, M.K. Samuelsson, A. Burchert, J.P. Bach, T.D. Kim, B. Berwanger, C. Thiede, R. Jager, G. Ehninger, H. Schafer, N. Ueki, M.J. Hayman, M. Eilers, and A. Neubauer. 2006. Inhibition of retinoic acid receptor signaling by Ski in acute myeloid leukemia. Leukemia 20:437–443.

Sahin, U., O. Ferhi, M. Jeanne, S. Benhenda, C. Berthier, F. Jollivet, M. Niwa-Kawakita, O. Faklaris, N. Setterblad, H. de The, and V. Lallemand-Breitenbach. 2014. Oxidative stress-induced assembly of PML nuclear bodies controls sumoylation of partner proteins. The Journal of cell biology 204:931–945.

Saitou, M., S. Sugai, T. Tanaka, K. Shimouchi, E. Fuchs, S. Narumiya, and A. Kakizuka. 1995. Inhibition of skin development by targeted expression of a dominant-negative retinoic acid receptor. Nature 374:159–162.

Shimomura, Y., H. Mitsui, Y. Yamashita, T. Kamae, A. Kanai, H. Matsui, T. Ishibashi, A. Tanimura, H. Shibayama, K. Oritani, J. Kuyama, and Y. Kanakura. 2016. New variant of acute promyelocytic leukemia with IRF2BP2-RARA fusion. Cancer Sci 107:1165–1168.

Sternsdorf, T., V.T. Phan, M.L. Maunakea, C.B. Ocampo, J. Sohal, A. Silletto, F. Galimi, M.M. Le Beau, R.M. Evans, and S.C. Kogan. 2006. Forced retinoic acid receptor alpha homodimers prime mice for APL-like leukemia. Cancer Cell 9:81–94.

Stielow, B., A. Sapetschnig, I. Kruger, N. Kunert, A. Brehm, M. Boutros, and G. Suske. 2008. Identification of SUMO-dependent chromatin-associated transcriptional repression components by a genome-wide RNAi screen. Mol Cell 29:742–754.

Tan, J., C.K. Ong, W.K. Lim, C.C. Ng, A.A. Thike, L.M. Ng, V. Rajasegaran, S.S. Myint, S. Nagarajan, S. Thangaraju, S. Dey, N.D. Nasir, G.C. Wijaya, J.Q. Lim, D. Huang, Z. Li, B.H. Wong, J.Y. Chan, J.R. McPherson, I. Cutcutache, G. Poore, S.T. Tay, W.J. Tan, T.C. Putti, B.S. Ahmad, P. Iau, C.W. Chan, A.P. Tang, W.S. Yong, P. Madhukumar, G.H. Ho, V.K. Tan, C.Y. Wong, M. Hartman, K.W. Ong, B.K. Tan, S.G. Rozen, P. Tan, P.H. Tan, and B.T. Teh. 2015. Genomic landscapes of breast fibroepithelial tumors. Nat Genet 47:1341–1345.

Tessier, S., O. Ferhi, M.C. Geoffroy, R. Gonzalez-Prieto, A. Canat, S. Quentin, M. Pla, M. Niwa-Kawakita, P. Bercier, D. Rerolle, M. Tirard, P. Therizols, E. Fabre, A.C.O. Vertegaal, H. de The, and V. Lallemand-Breitenbach. 2022. Exploration of nuclear body-enhanced sumoylation reveals that PML represses 2-cell features of embryonic stem cells. Nat Commun 13:5726.

Tharuka, M.D.N., A.S. Courelli, and Y. Chen. 2025. Immune regulation by the SUMO family. Nat Rev Immunol 25:608–620.

Theurillat, I., I.A. Hendriks, J.C. Cossec, A. Andrieux, M.L. Nielsen, and A. Dejean. 2020. Extensive SUMO Modification of Repressive Chromatin Factors Distinguishes Pluripotent from Somatic Cells. Cell Rep 33:108251.

Tiefenbach, J., N. Novac, M. Ducasse, M. Eck, F. Melchior, and T. Heinzel. 2006. SUMOylation of the corepressor N-CoR modulates its capacity to repress transcription. Mol Biol Cell 17:1643–1651.

Tsai, S., S. Bartelmez, R. Heyman, K. Damm, R. Evans, and S.J. Collins. 1992. A mutated retinoic acid receptor-alpha exhibiting dominant-negative activity alters the lineage development of a multipotent hematopoietic cell line. Genes Dev 6:2258–2269.

Tsuzuki, S., K. Kitajima, T. Nakano, A. Glasow, A. Zelent, and T. Enver. 2004. Cross talk between retinoic acid signaling and transcription factor GATA-2. Mol Cell Biol 24:6824–6836.

Valima, E., V. Varis, K. Bureiko, J.K. Lempiainen, A.M. Schroderus, L. Oksa, O. Lohi, T. Kinnunen, M. Varjosalo, E.A. Niskanen, V. Paakinaho, and J.J. Palvimo. 2025. SUMOylation inhibition potentiates the glucocorticoid receptor to program growth arrest of acute lymphoblastic leukemia cells. Oncogene 44:1259–1271.

Voisset, E., E. Moravcsik, E.W. Stratford, A. Jaye, C.J. Palgrave, R.K. Hills, P. Salomoni, S.C. Kogan, E. Solomon, and D. Grimwade. 2018. Pml nuclear body disruption cooperates in APL pathogenesis and impairs DNA damage repair pathways in mice. Blood 131:636–648.

Wang, P., S. Benhenda, H. Wu, V. Lallemand-Breitenbach, T. Zhen, F. Jollivet, L. Peres, Y. Li, S.J. Chen, Z. Chen, H. de The, and G. Meng. 2018. RING tetramerization is required for nuclear body biogenesis and PML sumoylation. Nat Commun 9:1277.

Westervelt, P., A.A. Lane, J.L. Pollock, K. Oldfather, M.S. Holt, D.B. Zimonjic, N.C. Popescu, J.F. DiPersio, and T.J. Ley. 2003. High-penetrance mouse model of acute promyelocytic leukemia with very low levels of PML-RARalpha expression. Blood 102:1857–1865.

Weston, A.D., B. Blumberg, and T.M. Underhill. 2003. Active repression by unliganded retinoid receptors in development: less is sometimes more. J Cell Biol 161:223–228.

Yin, C.C., N. Jain, M. Mehrotra, J. Zhagn, A. Protopopov, Z. Zuo, N. Pemmaraju, C. DiNardo, C. Hirsch-Ginsberg, S.A. Wang, L.J. Medeiros, L. Chin, K.P. Patel, F. Ravandi, A. Futreal, and C.E. Bueso-Ramos. 2015. Identification of a novel fusion gene, IRF2BP2-RARA, in acute promyelocytic leukemia. J Natl Compr Canc Netw 13:19–22.

Zhang, A., and S. Qiu. 2025. Advances in RARalpha fusion genes in acute promyelocytic leukemia. Exp Hematol 149:104822.

Zhang, L., X. Wang, D. Hu, S. Li, M. Sun, Q. Liu, H. Feng, M. Zhou, C. Chen, H. Zhou, and S. Ma. 2025. SUMOylation facilitates the stability of BCR-ABL to promote chronic myeloid leukemia progression. Oncogene 44:1844–1855.

Zhu, J., J. Zhou, L. Peres, F. Riaucoux, N. Honore, S. Kogan, and H. de The. 2005. A sumoylation site in PML/RARA is essential for leukemic transformation. Cancer Cell 7:143–153.

