## supplementary material for "PML-driven sumoylation of PML/RARA-bound co-repressors drives immortalization of primary hematopoietic progenitors"

Supplemental fig. 1

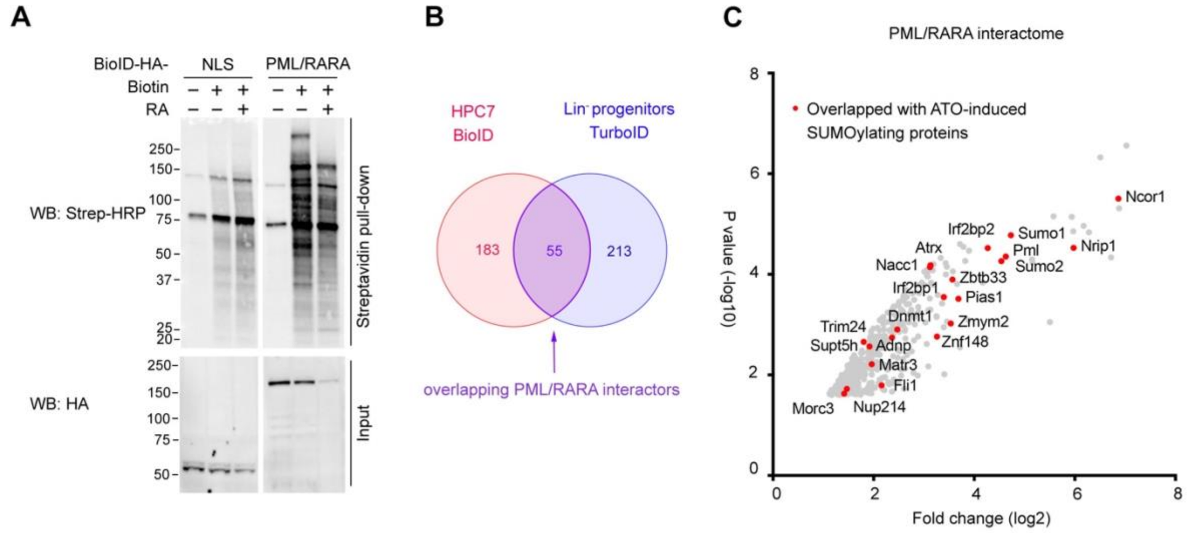


Fig. S1. **PML/RARA interacting proteins.** (**A**) Western blot analysis of lysate and streptavidin-purified fractions from HPC cell lines expressing NLS-BioID or PML/RARA-BioID with or without 24h all-trans-retinoic acid (ATRA) treatment. (**B**) Venn diagram showing overlap between PML/RARA interactors identified by BioID in HPCs and TurboID in PML/RARA-transduced Lin$^{-}$ progenitors. (**C**) Overlap of PML/RARA interactors in HPC7 cells with 1h arsenic (ATO)-induced SUMOylated proteins in primary mouse APL blasts.

Supplemental fig. 2

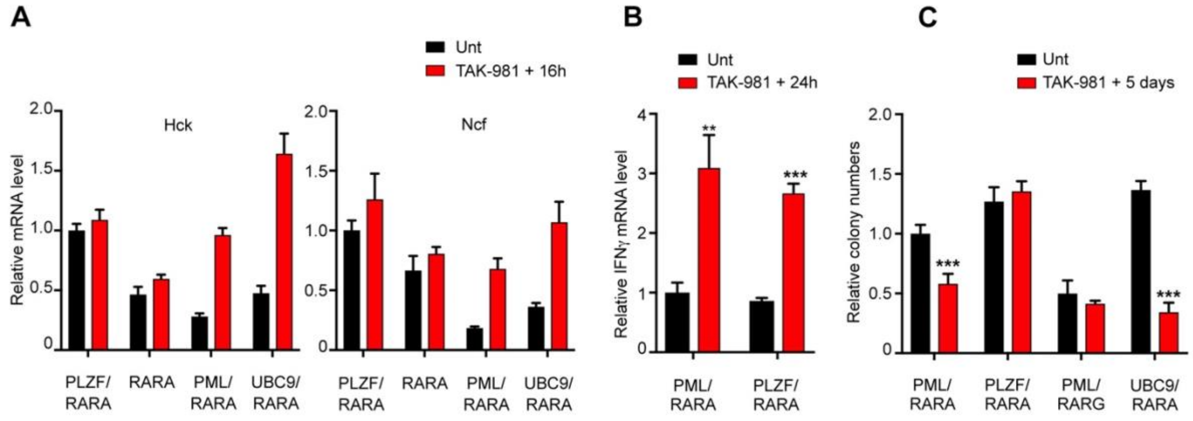


Fig S2. **TAK-981 restores RARA target gene expression to promote differentiation and impair colony formation.** (**A**) Real-time PCR for RARA targets, *Hck* and *Ncf*, gene expression in primary progenitors transformed with indicated constructs post TAK-981 (100 nM) treatment for 16h. (**B**) Real-time PCR performed in triplicate for *Ifng* expression in PML/RARA or PLZF/RARA transformed progenitors treated with TAK-981 (100 nM) for 24h. (**A**, **B**, and **C**) Data represent mean ± SD of triplicates; unpaired t test: $^{*}p < 0.05$, $^{**}p < 0.005$, $^{***}p < 0.001$; representative of three independent experiments.

**Supplementary Table 1.** List of 238 significant PML/RARA associated protein identified in HPC7 cell lines.

| Rank | Genes | Fold change (Biotin_vs_NLS) | P_Value (Biotin_vs_NLS) | FDR |
|---|---|---|---|---|
| 1 | Ncor2 | 228,83 | 1,02549E-07 | 1% |
| 2 | Ncor1 | 128,31 | 4,03815E-07 | 1% |
| 3 | Ski | 101,57 | 8,84751E-07 | 1% |
| 4 | Nrip1 | 93,06 | 1,17931E-06 | 1% |
| 5 | Rara | 90,99 | 8,34082E-06 | 1% |
| 6 | Ncoa3 | 67,02 | 3,13295E-06 | 1% |
| 7 | Jmjd1c | 62,33 | 2,51586E-06 | 1% |
| 8 | Gps2 | 54,00 | 3,45844E-06 | 1% |
| 9 | Sumo2;Sumo3 | 47,59 | 3,40478E-05 | 1% |
| 10 | Rxra | 41,14 | 2,47595E-06 | 1% |
| 11 | Rarb;Rarg | 39,15 | 0,00027642 | 1% |
| 12 | Ncoa2 | 37,52 | 8,58931E-06 | 1% |
| 13 | Hivep2 | 30,64 | 1,70207E-05 | 1% |
| 14 | Pml | 25,56 | 1,26128E-05 | 1% |
| 15 | Sumo1 | 22,90 | 9,03762E-06 | 1% |
| 16 | Irf2bp2 | 21,96 | 1,31805E-05 | 1% |
| 17 | Dkc1 | 16,61 | 1,38699E-05 | 1% |
| 18 | Foxk2 | 15,55 | 7,89158E-05 | 1% |
| 19 | Pias2 | 14,34 | 0,000305946 | 1% |
| 20 | Asxl2 | 12,59 | 2,26072E-05 | 1% |
| 21 | Rxrb | 11,98 | 0,000324268 | 1% |
| 22 | Mef2d | 11,74 | 0,000250719 | 1% |
| 23 | Cic | 10,66 | 6,09817E-05 | 1% |
| 24 | Nr2c2 | 10,07 | 0,000890133 | 1% |
| 25 | Cyb5r3 | 8,48 | 0,000138084 | 1% |
| 26 | Etv6 | 8,19 | 0,000175948 | 1% |
| 27 | Med1 | 7,71 | 0,005019948 | 5% |
| 28 | Tet2 | 7,40 | 0,000316152 | 1% |
| 29 | Nacc1 | 6,63 | 0,001652715 | 1% |
| 30 | Svil | 6,48 | 0,000514406 | 1% |
| 31 | Letm1 | 6,26 | 0,000462075 | 1% |
| 32 | Tmem70 | 6,12 | 0,0015988 | 1% |
| 33 | B2m | 6,01 | 0,004678042 | 5% |
| 34 | Cdipt | 5,94 | 0,000749746 | 1% |
| 35 | Slc25a12 | 5,67 | 0,010687363 | 5% |

| | | | | |
|---|---|---|---|---|
| 36 | Nipsnap1 | 5,60 | 0,002666425 | 5% |
| 37 | Acadvl | 5,49 | 0,002685824 | 5% |
| 38 | PML-RARA | 5,43 | 0,008920433 | 5% |
| 39 | Nxf1 | 5,36 | 0,001786744 | 1% |
| 40 | Fli1 | 5,32 | 0,003807212 | 5% |
| 41 | Foxk1 | 5,21 | 0,003317795 | 5% |
| 42 | Timm44 | 5,15 | 0,00825333 | 5% |
| 43 | Erg | 5,13 | 0,004742256 | 5% |
| 44 | Abcb10 | 5,08 | 0,006044575 | 5% |
| 45 | Ctbp2 | 5,05 | 0,003900758 | 5% |
| 46 | Arid1b | 4,88 | 0,005600205 | 5% |
| 47 | Nt5dc2 | 4,85 | 0,011427664 | 5% |
| 48 | Rab27a | 4,85 | 0,000789427 | 1% |
| 49 | Psmc5 | 4,82 | 0,004531378 | 5% |
| 50 | Lrpap1 | 4,56 | 0,002585669 | 5% |
| 51 | Dapp1 | 4,51 | 0,010200578 | 5% |
| 52 | Cct3 | 4,50 | 0,001699806 | 1% |
| 53 | Naxd | 4,50 | 0,012117956 | 5% |
| 54 | Fen1 | 4,49 | 0,005982131 | 5% |
| 55 | Mtnd1 | 4,45 | 0,012732733 | 5% |
| 56 | Dvl1 | 4,37 | 0,002017186 | 5% |
| 57 | Vps29 | 4,35 | 0,00135758 | 1% |
| 58 | Slc25a5 | 4,34 | 0,012091649 | 5% |
| 59 | Bak1 | 4,22 | 0,007931369 | 5% |
| 60 | Nnt | 4,22 | 0,008269747 | 5% |
| 61 | Cfap20 | 4,21 | 0,00281672 | 5% |
| 62 | Cs | 4,20 | 0,006745657 | 5% |
| 63 | Rab2a | 4,19 | 0,003548107 | 5% |
| 64 | Slc25a4 | 4,17 | 0,022382236 | 5% |
| 65 | Napsa | 4,12 | 0,003808393 | 5% |
| 66 | Icam2 | 4,12 | 0,006145137 | 5% |
| 67 | Myo1g | 4,12 | 0,011471846 | 5% |
| 68 | Fbl | 4,07 | 0,017335408 | 5% |
| 69 | Rac1 | 4,04 | 0,013427499 | 5% |
| 70 | Trappc5 | 4,02 | 0,003182474 | 5% |
| 71 | Ero1a | 4,00 | 0,013478821 | 5% |
| 72 | Pde12 | 3,97 | 0,005866062 | 5% |
| 73 | Psmd14 | 3,95 | 0,014841121 | 5% |
| 74 | Polr2b | 3,94 | 0,004750812 | 5% |
| 75 | Pdk3 | 3,94 | 0,004167981 | 5% |
| 76 | Sar1a;Sar1b | 3,93 | 0,003603422 | 5% |
| 77 | Trmt10c | 3,91 | 0,003377636 | 5% |
| 78 | Atp5pb | 3,88 | 0,005448294 | 5% |

| | | | | |
|---|---|---|---|---|
| 79 | Ctbp1 | 3,86 | 0,00501599 | 5% |
| 80 | Slc25a24 | 3,83 | 0,019921942 | 5% |
| 81 | Ykt6 | 3,81 | 0,011104951 | 5% |
| 82 | Arl1 | 3,76 | 0,012524012 | 5% |
| 83 | Dhx9 | 3,75 | 0,008365871 | 5% |
| 84 | Hdac1 | 3,74 | 0,005341223 | 5% |
| 85 | Eif3b | 3,71 | 0,014620218 | 5% |
| 86 | Cstf3 | 3,70 | 0,007416556 | 5% |
| 87 | Ndufs2 | 3,69 | 0,005641276 | 5% |
| 88 | Adnp | 3,68 | 0,00292408 | 5% |
| 89 | Arpc4 | 3,66 | 0,005107278 | 5% |
| 90 | Psmd1 | 3,65 | 0,004707997 | 5% |
| 91 | Syncrip | 3,62 | 0,00617159 | 5% |
| 92 | Rab5c | 3,58 | 0,005503934 | 5% |
| 93 | Mrpl38 | 3,55 | 0,007422696 | 5% |
| 94 | Lrpprc | 3,54 | 0,019832571 | 5% |
| 95 | Mtco2 | 3,54 | 0,011839196 | 5% |
| 96 | Rab8b | 3,54 | 0,006348106 | 5% |
| 97 | Eftud2 | 3,53 | 0,018427249 | 5% |
| 98 | Acad9 | 3,53 | 0,012747644 | 5% |
| 99 | Ptbp1 | 3,52 | 0,016664887 | 5% |
| 100 | Abcb7 | 3,52 | 0,010984513 | 5% |
| 101 | Gnpat | 3,51 | 0,009511217 | 5% |
| 102 | Otub1 | 3,51 | 0,006142601 | 5% |
| 103 | Pcbp2 | 3,51 | 0,00935894 | 5% |
| 104 | Dnajc8 | 3,50 | 0,011637808 | 5% |
| 105 | Eci1 | 3,49 | 0,003979555 | 5% |
| 106 | Cdk6 | 3,49 | 0,006834703 | 5% |
| 107 | Gatad2a | 3,48 | 0,020692243 | 5% |
| 108 | Nipsnap2 | 3,47 | 0,01587127 | 5% |
| 109 | Rcc2 | 3,46 | 0,009887154 | 5% |
| 110 | Ganab | 3,45 | 0,005606174 | 5% |
| 111 | Dars1 | 3,43 | 0,012195444 | 5% |
| 112 | Slc25a20 | 3,43 | 0,007546792 | 5% |
| 113 | Rpl23 | 3,42 | 0,018824296 | 5% |
| 114 | Lin54 | 3,41 | 0,02297903 | 5% |
| 115 | Cyp51a1 | 3,39 | 0,007292009 | 5% |
| 116 | Mob2 | 3,37 | 0,010022845 | 5% |
| 117 | Dpp3 | 3,35 | 0,006184357 | 5% |
| 118 | Ndufs3 | 3,34 | 0,012798494 | 5% |
| 119 | Iscu | 3,34 | 0,006530916 | 5% |
| 120 | Pccb | 3,34 | 0,013566969 | 5% |
| 121 | Sf3b3 | 3,34 | 0,00505165 | 5% |

| | | | | |
|---|---|---|---|---|
| 122 | Msh6 | 3,33 | 0,018019671 | 5% |
| 123 | Mov10 | 3,31 | 0,005178362 | 5% |
| 124 | Lnpep | 3,31 | 0,005492699 | 5% |
| 125 | Prkacb | 3,30 | 0,009257369 | 5% |
| 126 | Prpf31 | 3,29 | 0,017602561 | 5% |
| 127 | Pdpr | 3,28 | 0,007685206 | 5% |
| 128 | Hells | 3,28 | 0,011071202 | 5% |
| 129 | Fkbp5 | 3,27 | 0,012858563 | 5% |
| 130 | Ezr | 3,27 | 0,005773631 | 5% |
| 131 | Aacs | 3,25 | 0,008913366 | 5% |
| 132 | Zhx2 | 3,24 | 0,009817101 | 5% |
| 133 | Pdcd10 | 3,24 | 0,011917044 | 5% |
| 134 | Mrps35 | 3,24 | 0,009437516 | 5% |
| 135 | Ndufa13 | 3,23 | 0,013280189 | 5% |
| 136 | Snw1 | 3,23 | 0,019523821 | 5% |
| 137 | Pmpcb | 3,23 | 0,024421634 | 5% |
| 138 | Idi1 | 3,23 | 0,015749777 | 5% |
| 139 | Sbno1;Sbno2 | 3,22 | 0,005585525 | 5% |
| 140 | Mcm3 | 3,22 | 0,023679927 | 5% |
| 141 | Dnajc11 | 3,21 | 0,005492505 | 5% |
| 142 | Lrrc59 | 3,20 | 0,01288013 | 5% |
| 143 | Nup93 | 3,18 | 0,006343833 | 5% |
| 144 | Ap2b1 | 3,17 | 0,011180123 | 5% |
| 145 | Sfxn1 | 3,17 | 0,013209045 | 5% |
| 146 | Acly | 3,16 | 0,013672404 | 5% |
| 147 | Atad3 | 3,16 | 0,015678194 | 5% |
| 148 | Ddx39a | 3,16 | 0,007011783 | 5% |
| 149 | Copa | 3,15 | 0,01421791 | 5% |
| 150 | Ddx19a | 3,15 | 0,009972919 | 5% |
| 151 | Eif4a3 | 3,14 | 0,01347579 | 5% |
| 152 | Uba2 | 3,13 | 0,014335835 | 5% |
| 153 | Slc44a2 | 3,13 | 0,023547122 | 5% |
| 154 | Gnas | 3,11 | 0,022752548 | 5% |
| 155 | Aldh9a1 | 3,11 | 0,023333018 | 5% |
| 156 | Phb2 | 3,11 | 0,018796223 | 5% |
| 157 | Rpl6 | 3,10 | 0,02022028 | 5% |
| 158 | Psmd12 | 3,10 | 0,013897792 | 5% |
| 159 | Ythdf2 | 3,09 | 0,00719557 | 5% |
| 160 | Bcat2 | 3,08 | 0,022170605 | 5% |
| 161 | Pdia3 | 3,06 | 0,016509834 | 5% |
| 162 | Snrnp200 | 3,06 | 0,0126063 | 5% |
| 163 | Tomm34 | 3,04 | 0,011612271 | 5% |
| 164 | Rab5a | 3,03 | 0,011388137 | 5% |

| | | | | |
|---|---|---|---|---|
| 165 | Vps4b | 3,02 | 0,016699134 | 5% |
| 166 | Tsn | 3,01 | 0,011597824 | 5% |
| 167 | Naa15 | 3,00 | 0,012354699 | 5% |
| 168 | Xpo7 | 2,99 | 0,013581436 | 5% |
| 169 | Mrps27 | 2,99 | 0,016262618 | 5% |
| 170 | Lgals9 | 2,99 | 0,019533785 | 5% |
| 171 | Slc25a1 | 2,99 | 0,016747008 | 5% |
| 172 | Fermt3 | 2,98 | 0,022716372 | 5% |
| 173 | Gsr | 2,97 | 0,018318922 | 5% |
| 174 | Vav1 | 2,97 | 0,022569345 | 5% |
| 175 | Paics | 2,96 | 0,008301189 | 5% |
| 176 | Mcm7 | 2,96 | 0,013835846 | 5% |
| 177 | Sdha | 2,95 | 0,024403465 | 5% |
| 178 | Epm2aip1 | 2,94 | 0,024596742 | 5% |
| 179 | Rfc3 | 2,93 | 0,024381241 | 5% |
| 180 | Sgpl1 | 2,93 | 0,013722637 | 5% |
| 181 | Rtn3 | 2,92 | 0,013148368 | 5% |
| 182 | Uggt1 | 2,91 | 0,014593034 | 5% |
| 183 | Eef1g | 2,89 | 0,012731617 | 5% |
| 184 | Nrm | 2,89 | 0,011643826 | 5% |
| 185 | Gna12;Gna13 | 2,88 | 0,012476454 | 5% |
| 186 | Hnrnpm | 2,88 | 0,017410011 | 5% |
| 187 | Rab33b | 2,88 | 0,015830231 | 5% |
| 188 | Nars2 | 2,87 | 0,022907362 | 5% |
| 189 | Snrpa1 | 2,87 | 0,011517675 | 5% |
| 190 | Spdl1 | 2,86 | 0,017731941 | 5% |
| 191 | Vdac3 | 2,86 | 0,014056957 | 5% |
| 192 | Mef2c | 2,86 | 0,018292759 | 5% |
| 193 | Supt5h | 2,86 | 0,018303309 | 5% |
| 194 | Rab6a | 2,85 | 0,018806153 | 5% |
| 195 | Rac2 | 2,85 | 0,019455569 | 5% |
| 196 | Prdx4 | 2,84 | 0,01743679 | 5% |
| 197 | Lyn | 2,83 | 0,019020669 | 5% |
| 198 | Ddx5 | 2,82 | 0,017488574 | 5% |
| 199 | Glo1 | 2,82 | 0,014882966 | 5% |
| 200 | Rpl12 | 2,82 | 0,022399137 | 5% |
| 201 | Cul3 | 2,81 | 0,012082237 | 5% |
| 202 | Eif4a1 | 2,81 | 0,012909909 | 5% |
| 203 | Ssrp1 | 2,79 | 0,02108522 | 5% |
| 204 | Dnajb1 | 2,77 | 0,020220632 | 5% |
| 205 | Arf4 | 2,75 | 0,015353245 | 5% |
| 206 | Stat5a;Stat5b | 2,75 | 0,016098393 | 5% |
| 207 | Lamp1 | 2,72 | 0,014420806 | 5% |

| | | | | |
|---|---|---|---|---|
| 208 | Drg1 | 2,72 | 0,017846848 | 5% |
| 209 | Asph | 2,72 | 0,020991603 | 5% |
| 210 | Rnf220 | 2,71 | 0,019305279 | 5% |
| 211 | Supt16h | 2,69 | 0,019806141 | 5% |
| 212 | Rab35 | 2,68 | 0,015932306 | 5% |
| 213 | Rab21 | 2,68 | 0,014234157 | 5% |
| 214 | Cpne1 | 2,68 | 0,016784301 | 5% |
| 215 | Eprs1 | 2,67 | 0,020879249 | 5% |
| 216 | Lypla2 | 2,67 | 0,01747809 | 5% |
| 217 | Sigmar1 | 2,66 | 0,024259212 | 5% |
| 218 | Arhgap1 | 2,65 | 0,021874664 | 5% |
| 219 | Sae1 | 2,64 | 0,019762746 | 5% |
| 220 | Hsdl1 | 2,64 | 0,024802032 | 5% |
| 221 | Rars1 | 2,63 | 0,022054499 | 5% |
| 222 | Tfrc | 2,63 | 0,015669017 | 5% |
| 223 | Smc1a | 2,61 | 0,024768821 | 5% |
| 224 | Hars2 | 2,60 | 0,019535271 | 5% |
| 225 | Atp5f1a | 2,60 | 0,020530255 | 5% |
| 226 | Hsd17b4 | 2,59 | 0,02049348 | 5% |
| 227 | Spen | 2,59 | 0,019708072 | 5% |
| 228 | Arhgap18 | 2,59 | 0,025015419 | 5% |
| 229 | Cdc23 | 2,58 | 0,024706527 | 5% |
| 230 | Tars1 | 2,58 | 0,022976335 | 5% |
| 231 | Prpf19 | 2,57 | 0,023992272 | 5% |
| 232 | Snd1 | 2,56 | 0,021190452 | 5% |
| 233 | Tmed7 | 2,55 | 0,022642912 | 5% |
| 234 | Eif3d | 2,54 | 0,02224298 | 5% |
| 235 | Nsf | 2,50 | 0,022675362 | 5% |
| 236 | Akr1c6 | 0,38 | 0,024100654 | 5% |
| 237 | S100a14 | 0,34 | 0,011453224 | 5% |
| 238 | Ckb | 0,25 | 0,006548406 | 5% |
